# Persistent vulnerability to relapse despite complete extinction of cocaine craving

**DOI:** 10.1101/050401

**Authors:** Paul Girardeau, Sylvia Navailles, Audrey Durand, Caroline Vouillac-Mendoza, Karine Guillem, Serge H Ahmed

## Abstract

Craving often precedes relapse into cocaine addiction. This explains why considerable research effort is being expended to try to develop anti-craving strategies for relapse prevention. Recently, we discovered using the classic reinstatement model of cocaine craving that the reinstating or priming effect of cocaine can be extinguished with repeated priming in rats - a phenomenon dubbed extinction of cocaine priming. Here we sought to measure the potential beneficial effect of this novel extinction strategy on subsequent relapse (i.e., return to the pre-extinction pattern of cocaine self-administration once the drug is made again available after extinction). Overall and contrary to our initial hope, extensive and complete extinction of cocaine priming had no major impact on relapse. This lack of effect occurred despite evidence for post-extinction loss of neuronal responses to cocaine priming in brain regions critically involved in cocaine-induced reinstatement (i.e., the dorsomedial prefrontal cortex and the core of the nucleus accumbens). An effect of extinction of cocaine priming on relapse was only observed when cocaine was available for self-administration under more demanding conditions. However, this effect was modest and short-lived. Finally, we succeeded to trace the origin of our failure to prevent relapse to a persistent, extinction-resistant form of operant behavior that is not directly induced by cocaine. This extinction-resistant behavior is commonly reported, though generally ignored as causally irrelevant, in many other reinstatement studies. We propose that this behavior should become both a novel marker for long-term vulnerability to relapse and a novel target for preclinical development of potential relapse prevention interventions.

---

Cocaine not only evokes intense pleasurable sensations but also induces an overwhelming desire or craving for more cocaine. This latter process is especially obvious in individuals with a diagnosis of cocaine use disorder and is thought to contribute, together with other factors, to precipitate or favor relapse after abstinence ^1, 2, 3^. Cocaine-induced craving can be approached and studied in animals using the classic drug reinstatement model ^4, 5^. In this model, operant responding for the drug (e.g., pressing a lever) is first extinguished by discontinuing drug reinforcement and then reinstated by drug priming (i.e., passive or non-contingent drug administration). Importantly, after drug priming, reinstated operant responding continues to be non-reinforced as during extinction and, therefore, is thought to be a behavioral expression of genuine drug seeking ^6, 7^. This standard model has been extensively used to study the neural correlates and substrates of cocaine seeking and to develop potential anti-craving interventions for relapse prevention ^8, 9, 10^.

Using this model, we recently discovered that the reinstating effect of cocaine on previously extinguished cocaine seeking can itself be extinguished with repeated drug priming - a phenomenon dubbed extinction of drug priming to distinguish it from extinction of operant drug responding ^11^. Such extinction has been interpreted as evidence that the priming effect of cocaine on reinstatement of cocaine seeking depends on an interoceptive drug conditioning mechanism whereby the interoceptive cues of cocaine become reliable conditioned Pavlovian predictors of the availability of cocaine reinforcement ^7, 11, 12, 13^. During cocaine priming, these cues would lead rats to expect, albeit falsely, that cocaine is again available for self-administration, thereby prompting them to reinstate responding. After repeated priming, however, these cues would progressively lose their predictive function. Regardless of the underlying mechanisms, however, extinction of drug priming has been proposed as a potential cocaine exposure therapy for relapse prevention that may complement other, more traditional exteroceptive cue exposure therapies ^11, 14^.

In theory, if the priming effect of cocaine played an important role in the vulnerability to relapse, as generally held, then one should expect that its extinction after repeated priming should prevent or at least retard relapse to cocaine self-administration ^11^. The overarching goal of the present series of experiments was to directly test this prediction. To achieve this end, we developed a within-session, multiple drug priming procedure (i.e., 3 per session) to promote a rapid and complete extinction of cocaine priming in rats with relatively long prior histories of intravenous cocaine self-administration (i.e., > 24 daily sessions) (see Methods). After extinction of cocaine priming, cocaine was again made available for self-administration under the same operant conditions as before extinction. Relapse was measured by comparing in the same individuals their pre-*versus* post-extinction patterns and rates of cocaine self-administration. To avoid confusion, the term relapse is strictly defined here as a return to the pre-extinction pattern and rate of cocaine self-administration - an operational definition that is close to the clinical definition ^4, 15^. In contrast and as already defined above, the term reinstatement is exclusively used to refer to the reinstatement of non-reinforced cocaine seeking by cocaine priming.

## Results

### Experiment 1: Effects of extinction of non-contingent cocaine priming on relapse

After extended cocaine self-administration training, extinction of cocaine priming was induced during 5 daily sessions of extinction (E1 to E5) (see Methods). Briefly, each session of extinction consisted of 3 cocaine primings (i.e., P1 to P3; 1 mg each, i.v.) administered 45 min apart following an initial 90-min period of extinction of operant responding. The effects of cocaine priming on reinstatement of operant responding was measured during the 45-min periods following P1, P2 and P3, and compared to non-primed responding (called NP) measured during the 45-min period preceding P1.

On session E1, all priming doses of cocaine (P1 to P3) reinstated responding above NP responding but this effect rapidly decreased within-session with repeated priming (i.e., from P1 to P3) (F3,57 = 27.57, p < 0.01) (Fig. 1a). A higher-resolution time course of the within-session extinction of cocaine priming is shown in Fig. S1. After repeated session of extinction, there was a further extinction of cocaine priming (Priming x Session: F3,57 = 17.58, p < 0.01). On session E5, all 3 priming doses of cocaine induced the same levels of reinstatement which were lower than those on session E1 (Fig. 1a). To better capture the extinction of cocaine priming across sessions, the effects of P1, P2 and P3 were averaged for each rat and each session. After an initially large extinction, the average priming effect of cocaine (i.e., P) rapidly levelled off at a residual level that was slightly, though significantly, above NP responding, suggesting that maximum extinction of cocaine priming was achieved (Priming x Session: F4,76 = 17.56, p < 0.01) (Fig. 1c). Finally, the extinction of cocaine priming was behaviorally-specific because no significant change in cocaine-induced increased locomotion was observed within-session or between-session (Fig. 1b, d).

**Figure 1.**
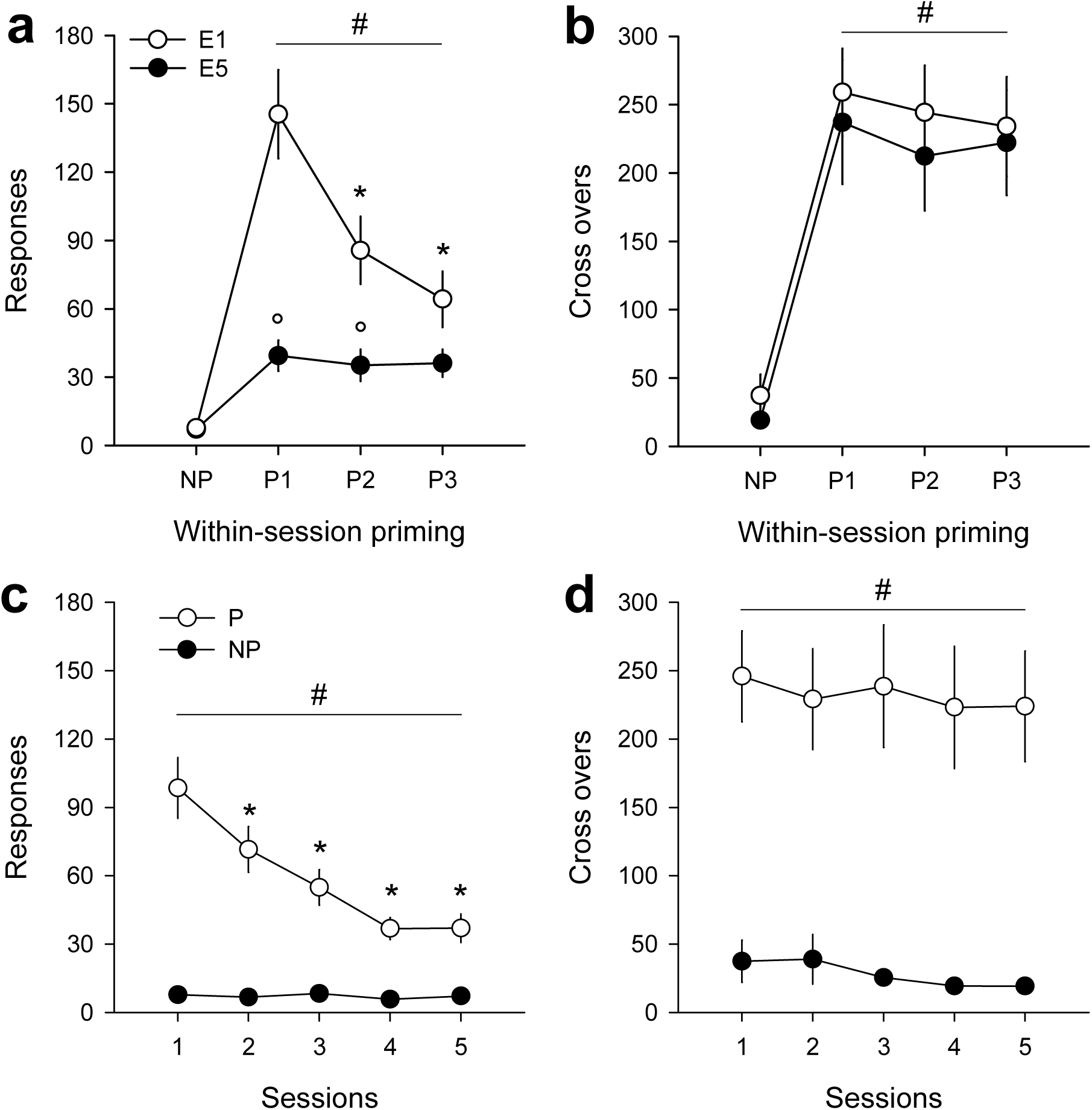
Extinction of cocaine priming. (**a**) Within-and between-session extinction of the effect of cocaine priming on non-reinforced responding (mean ± s.e.m.). After an initial non-primed period of 90 min, rats received intravenously 3 non-contingent priming doses of 1-mg cocaine (P1 to P3) spaced 45 min apart. The effect of each priming dose of cocaine was compared to non-primed responding (NP) measured during the 45-min period preceding P1. #, different from NP during the first (E1) and last session of extinction (E5); *, different from P1; °, different from E1 (p < 0.01). (**b**) Cocaine-induced increased cage crossovers (mean ± s.e.m.) during extinction testing #, different from NP during both the first (E1) and last session of extinction (E5) (p < 0.01). (**c**) Extinction of the average effect of cocaine priming (i.e., P = average of P1, P2 and P3) across sessions (mean ± s.e.m.). #, different from NP; *, different from session 1 (p < 0.01). (**d**) No change in the average effect of cocaine priming on cage crossovers during testing (mean ± s.e.m.). #, different from NP (p < 0.01).

On the day after extinction of cocaine priming, rats were given again access to cocaine for self-administration as before extinction to measure relapse. Overall, no effect on relapse to cocaine self-administration was observed. The pre-versus post-extinction patterns of cocaine self-administration were nearly identical (Fig. 2a). Though some rats were slightly slower to reinitiate cocaine self-administration within the first post-extinction session, they nevertheless readily re-expressed the same pattern of self-administration as before extinction once it was initiated. This was confirmed by a quantitative analysis of the rate of cocaine self-administration (i.e., total number of injections in the session divided by the time elapsed since initiation of cocaine self-administration). Clearly, after extinction of cocaine priming, rats took cocaine at a rate identical to or slightly higher than pre-extinction (average of the last 3 pre-extinction sessions), from post-extinction session 1 onward (F5,95 = 7.31, p < 0.01) (Fig. 2b).

**Figure 2.**
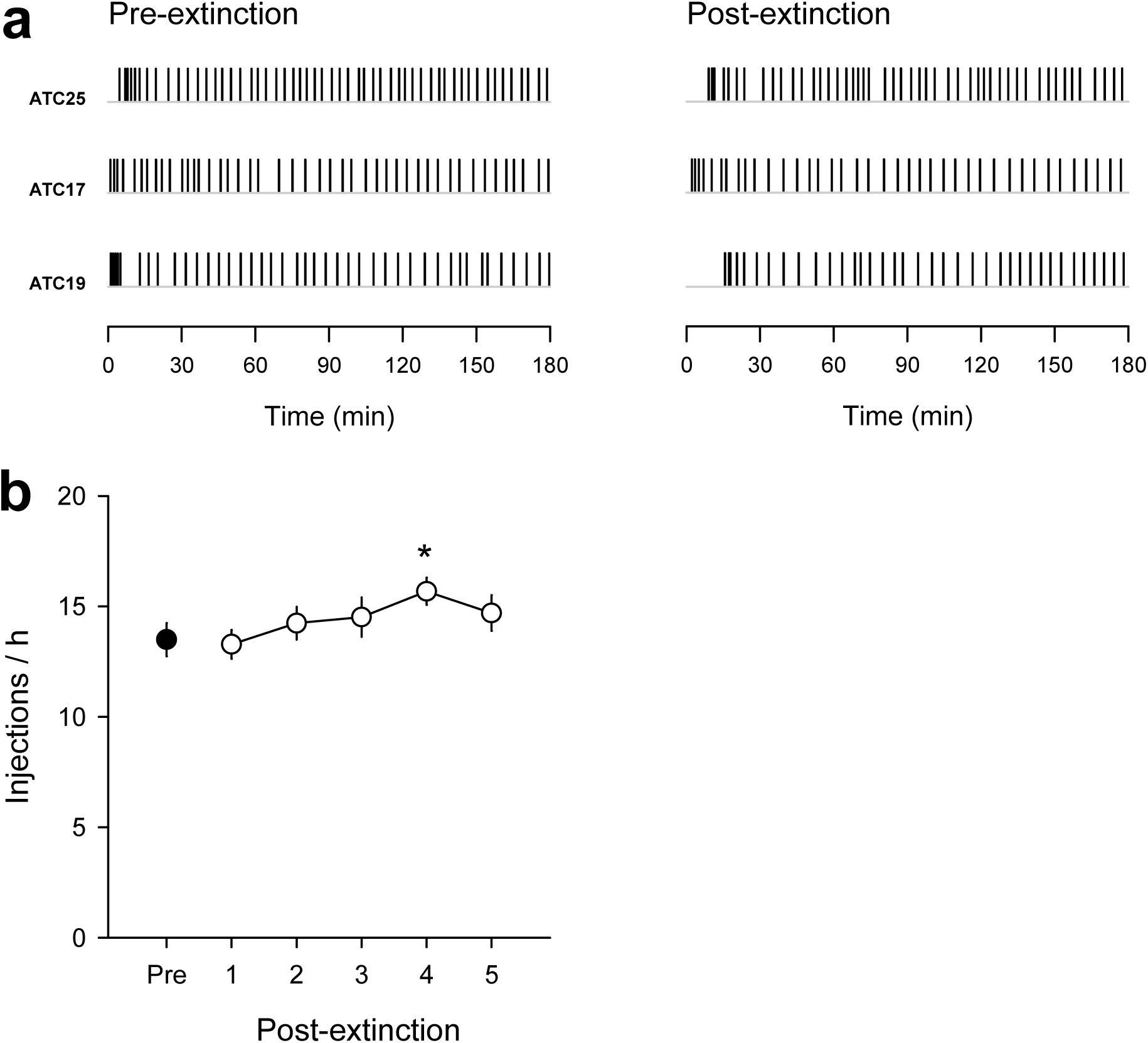
Post-extinction relapse to cocaine self-administration. (**a**) Pre-versus post-extinction temporal patterns of cocaine injections in 3 representative rats. Pre-extinction patterns were obtained on the last session of self-administration preceding extinction. Post-extinction patterns were obtained on the first session of relapse after extinction. Each vertical tick represents an i.v. injection of 0.25-mg cocaine under a FR3 procedure of self-administration. (**b**) Pre-versus post-extinction rate of cocaine self-administration (mean ± s.e.m.). The pre-extinction rate was obtained by averaging the last 3 sessions of self-administration that preceded extinction of cocaine priming. *, different from pre-extinction (p < 0.01).

### Experiment 2: Effects of extinction of response-contingent cocaine priming on relapse

The absence of relapse prevention despite extinction of cocaine priming may be related to the passive, involuntary nature of our multiple-priming procedure. To test this, a separate group of rats was given voluntary control over cocaine priming. Briefly, after an initial 45-min period of extinction, each priming period (P1 to P3), including the NP period preceding P1 was initiated if rats completed the operant response requirement within 25 min (see Methods for additional information). On 28 rats tested, 8 rats failed to obtain at least one self-priming on every session and were thus excluded from analysis. The remaining 20 rats obtained on average 2.7 ± 0.1 self-primings per session which was close to the maximum possible. However, though rats self-initiated several NP periods (4.9 ± 0.3 out of a maximum of 6), they failed to self-initiate one NP period on every session of extinction. Consequently, we could not compare the average priming effect of cocaine to NP responding on every session of extinction. To circumvent this limitation, we averaged for each rat its level of NP responding across all the sessions where it self-initiated a NP period. This should not be a major limitation, however; as shown in the other experiments of the present study, the level of NP responding was generally stable across sessions.

As with non-contingent cocaine priming, self-priming effectively reinstated responding above NP responding (session 1 to 5: F1,19 > 18.00, p < 0.01; session 6: F1,19 = 5.93, p < 0.05) and this effect decreased considerably, though not completely, with repeated session (F5,95 = 31.59, p < 0.01) (Fig. 3a). This between-session extinction of cocaine self-priming was not associated with a change in cocaine-induced increased locomotion (F5,95 = 0.58) (Fig. 3b). Finally, despite extinction of cocaine self-priming, there was no significant impact on relapse to cocaine self-administration post-extinction (F6,14 = 1.26) (Fig. 3c,d). The rate of cocaine self-administration returned to the pre-extinction level as early as the first post-extinction session onwards.

**Figure 3.**
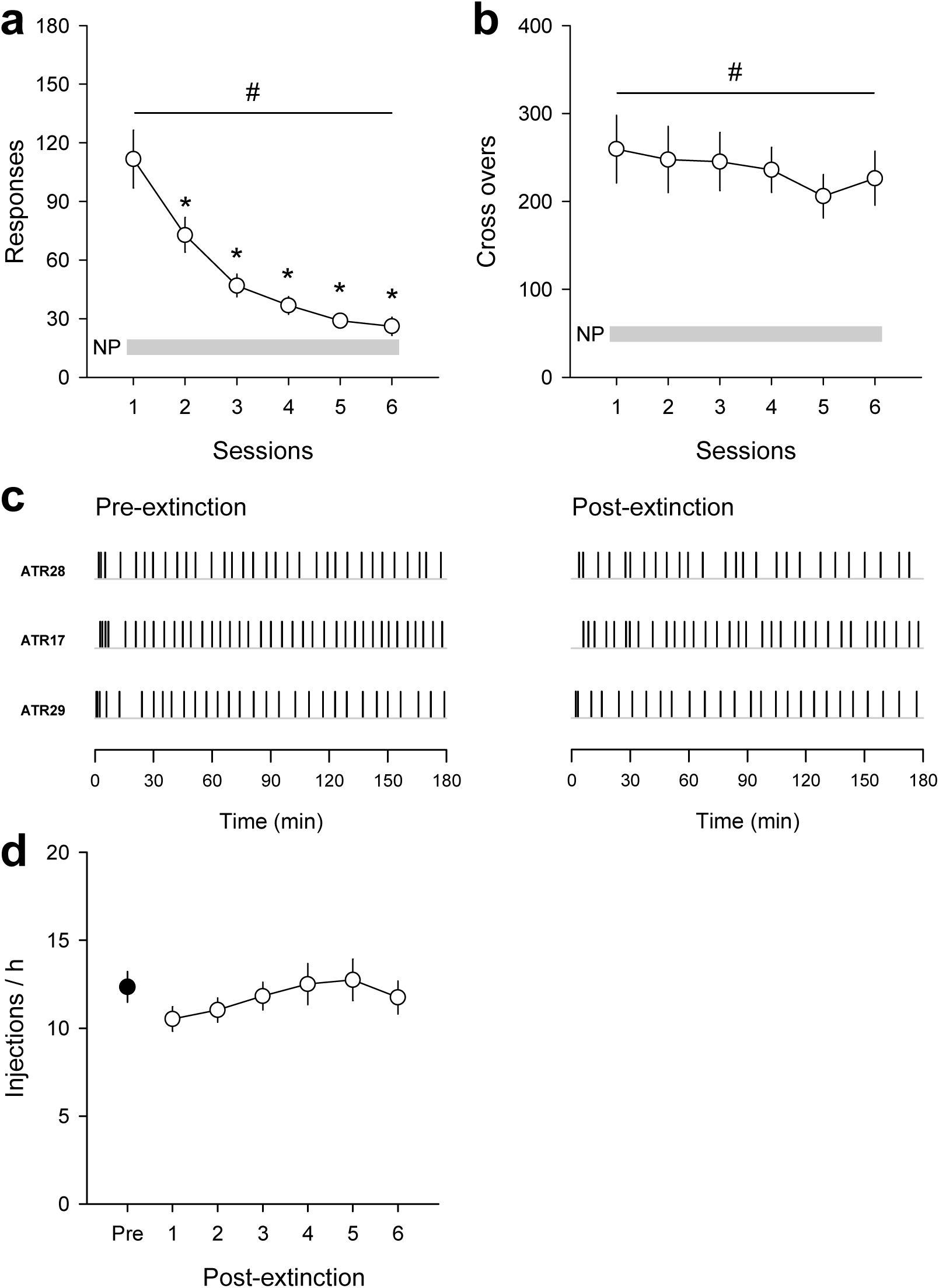
Relapse to cocaine self-administration after extinction of cocaine self-priming. (**a**) Extinction of the average effect of cocaine self-priming across sessions of extinction (mean ± s.e.m.). #, different from NP; *, different from session 1 (p < 0.01). (**b**) No change in the average stimulant effect of cocaine self-priming on cage crossovers during testing (mean ± s.e.m.). #, different from NP (p < 0.01). (**c**) Pre-versus post-extinction within-session patterns of cocaine injections in 3 representative rats. (**d**) Pre-versus post-extinction rate of cocaine self-administration (mean ± s.e.m.) (see Figure 2 for additional information).

### Experiment 3: Effects of extensive extinction of cocaine priming on relapse

In the two previous experiments, despite a large extinction of cocaine priming, there remained a small residual effect of cocaine priming that could be sufficient to explain post-extinction relapse to cocaine self-administration. To test if this residue could eventually be totally extinguished with additional sessions of extinction to prevent relapse, another group of rats were tested for a total of 24 sessions. These sessions were organized in 4 extinction blocks of 6 sessions each that alternated with 4 relapse blocks of 6 sessions of cocaine self-administration. This was done to follow any change in vulnerability to relapse with increased sessions of extinction. There was clear evidence for a large extinction of cocaine priming with repeated sessions and across blocks of extinction (Priming x Session: F23,391 = 3.74, p < 0.01) (Fig. 4a). Interestingly, there was some evidence for reconditioning of cocaine priming between extinction blocks (Session [1 versus 6] x Block: F3,51 = 7.17, p < 0.01) (Fig. 4a, b). Specifically, the effect of cocaine priming on session 1 of extinction block 2 increased above that of session 6 of block 1 (Fig. 4b). However, this reconditioning effect was short-lived. It was not observed during the following blocks of extinction (Fig. 4b), indicating a stable retention of extinction of cocaine priming across blocks. On the last 3 sessions of the final block of extinction, the priming effect of cocaine was very low compared to the first session of block 1 (F3,51 = 9.78, p < 0.01) but was still significantly above NP responding (Fig. 4a,c). Overall, there was no effect on relapse to cocaine self-administration between blocks of extinction (F24,408 = 1.44) (Fig. 4d) (see Fig. S2 for representative pre-and post-extinction patterns of cocaine self-administration).

**Figure 4.**
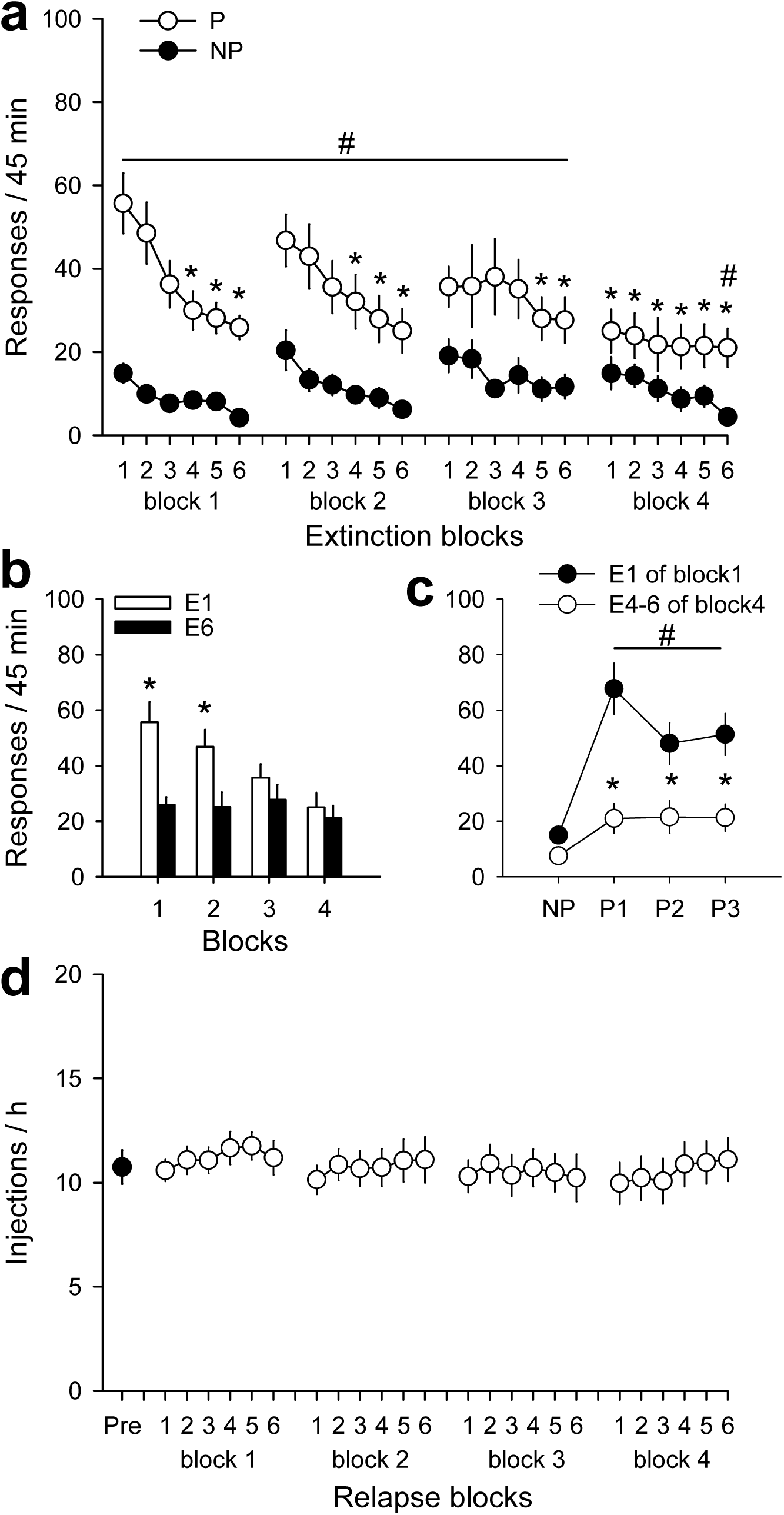
Relapse to cocaine self-administration after extensive extinction of cocaine priming. (**a**) Extinction of the average effect of cocaine priming across 4 extinction blocks of 6 sessions (E1-6) (mean ± s.e.m.). Blocks of extinction alternated with blocks of 6 relapse sessions. #, different from NP; *, different from session 1 (p < 0.01). (**b**) Average effect of cocaine priming during the first (E1) versus last (E6) session of extinction across blocks. *, different from all the last sessions (i.e., E6) of all blocks (p < 0.01). (**c**) Extinction of the effect of all 3 priming doses of cocaine across sessions (see Figure 1 for additional information). #, different from NP during the first session of block 1 (E1) and the last 3 stable sessions of block 4 (E4-6); *, different from E1 (p < 0.01). (**d**) Pre-versus post-extinction rate of cocaine self-administration across the 4 relapse blocks that alternated with the 4 extinction blocks (mean ± s.e.m.).

### Experiment 4: Effects of saline substitution on residual cocaine priming after extinction

Apparently, the priming effect of cocaine cannot be entirely extinguished, even after extensive extinction testing. This prompted us to investigate the underlying mechanisms of the post-extinction residual effect of cocaine priming. We first checked whether this effect was attributable to the intravenous infusion procedure itself. This was particularly warranted in the present study because no intravenous infusion was administered during the NP period preceding P1. To test this possibility, after extinction of cocaine priming and evidence for a residual effect of cocaine priming, cocaine was replaced with saline - its vehicle - all else being equal. Saline substitution was replicated 3 times between regular sessions of cocaine extinction. Overall, no residual effect was observed after saline substitution (F3,36 = 14.57, p < 0.01) (Fig. S3a), confirming that this effect was due to cocaine priming and not to the intravenous infusion procedure.

### Experiment 5: Effects of light cue conditions on residual cocaine priming after extinction

We next tested an eventual role for the response-contingent light cue above the operant lever in the residual effect of cocaine priming. To this end, rats from experiment 3 were tested during 21 additional sessions of extinction under 3 light cue conditions: light cue present (LC); no light cue available (NLC); and house light cue present (HLC) (see Methods).

Overall, there was a main effect of sessions, indicating an influence of the light cue condition on the residual effect of cocaine priming (F20,340 = 2.6, p < 0.01) (Fig. 5a). Subsequent analysis of data averaged across all sessions with the same light cue condition revealed that removal of the light cue (NLC condition) above the operant lever abolished the residual effect of cocaine priming compared to the LC condition (F2,34 = 4.33 p < 0.05) (Fig. 5b). Importantly, this effect was probably due to some unconditioned visual properties of the light cue because the residual effect of cocaine re-emerged in presence of a novel response-contingent light cue never associated with cocaine (i.e., a diffuse light from the cage roof, HLC) (Fig. 5b).

**Figure 5.**
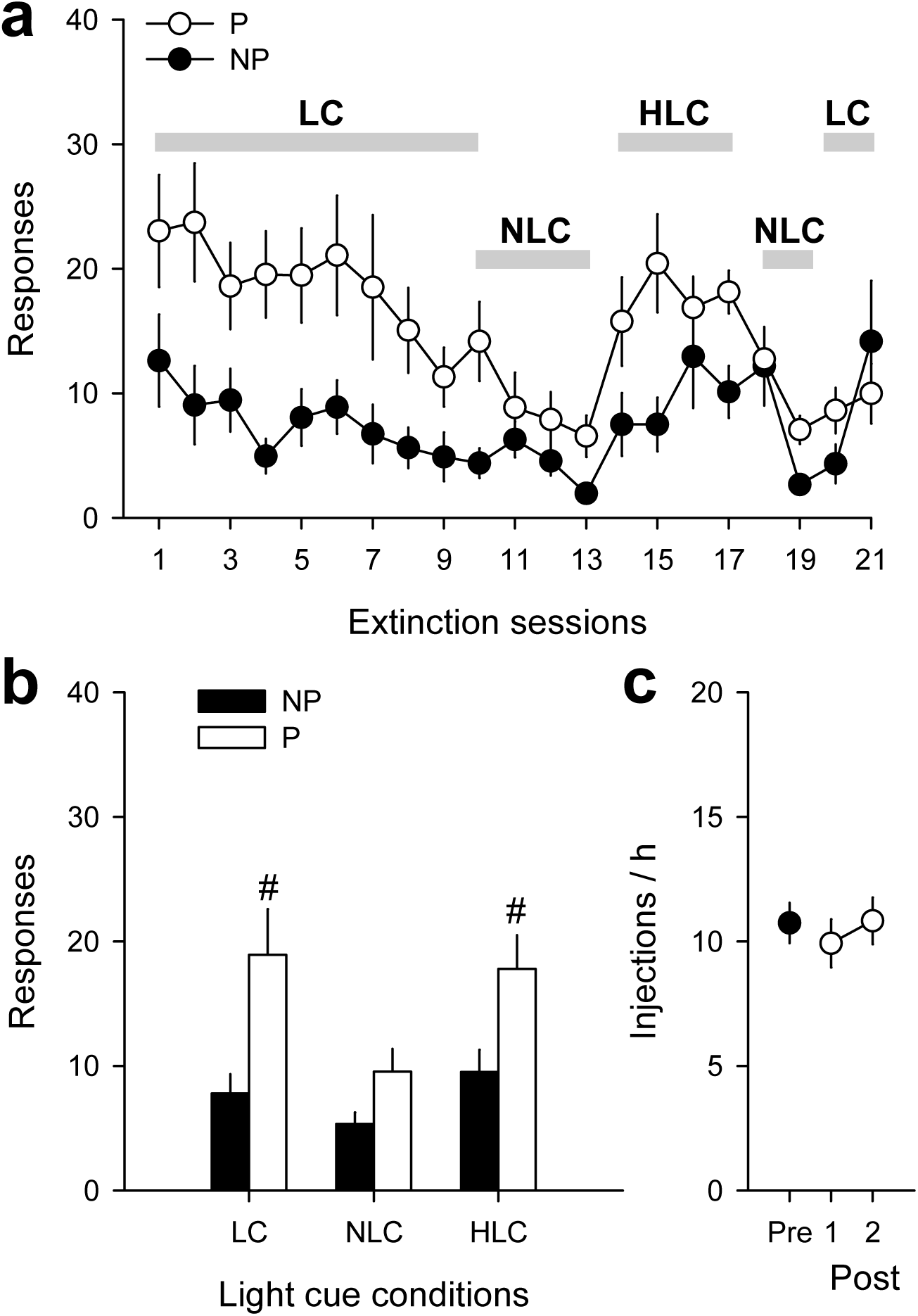
Cocaine potentiation of operant responding for a light cue. (**a**) Effects of different light cue conditions on the average effect of cocaine priming. The following three light conditions were tested for at least 4 sessions each (indicated by an horizontal grey bar above the curve): LC, response-contingent presentation of the light cue previously associated with cocaine; NLC, no response-contingent light cue is presented; HLC, response-contingent presentation of the house light cue. (**b**) Average effects across all testing sessions of the different light cue conditions. *, different from NP (p < 0.01). (**c**) Pre-versus post-extinction rate of cocaine self-administration (mean ± s.e.m.).

### Experiment 6: Effects of cocaine on operant responding for a neutral light cue after extinction of cocaine priming

The effect of the HLC condition was further confirmed in an independent group of rats that were trained to self-administer cocaine self-administration with no light cue (Fig. S4).

### Experiment 7: Effects of cocaine on operant responding for a neutral light cue in naïve rats

To rule out an eventual stimulus generalization effect, we tested the effects of cocaine on operant responding for a neutral light cue in a group of initially drug-and operant-naïve rats (n = 28). During testing, they received increasing i.v. doses of cocaine during access to an operant lever that turned on a light cue above it for 20 s under a FR1 procedure. Cocaine dose-dependently increased responding when a response-contingent light cue was present but not when it was absent (Dose x Cue: F3,81 = 4.23, p < 0.01) (Fig. S3b). The latter findings demonstrate that the residual effect of cocaine priming observed previously does not reflect a residue of cocaine seeking but a drug potentiation of some unconditioned reinforcing effects of light itself, whatever they may be.

### Experiment 8: Effects of extensive and complete extinction of cocaine priming on relapse

In experiments 3 and 5, the same rats (n = 18) received a total of 45 sessions of extinction of cocaine priming. Interestingly, during the last two sessions of extinction, there was no longer evidence for a residual effect of cocaine priming, suggesting its complete extinction (F1,17 = 0.00) (Fig. 5a). We took advantage of this to measure the effect on relapse to cocaine for self-administration. Despite extensive and complete extinction of cocaine priming, however, there was still no significant impact on relapse (F2,34 = 1.30) (Fig. 5c) (see Fig. S2 for representative pre-and post-extinction patterns of cocaine self-administration).

### Experiment 9: Effects of extinction of cocaine priming on relapse under a more demanding procedure of cocaine self-administration

In all previous experiments, a relatively low level of effort (i.e., FR3) was required to obtain cocaine during both maintenance of and relapse to drug self-administration. This factor may contribute to explain why extinction of cocaine priming had no major impact on relapse. To test this hypothesis, rats were trained to self-administer cocaine under a more demanding FR procedure. This procedure also allowed measurement of several behavioral parameters reflecting the motivation for the drug ^16, 17, 18^. Briefly, the FR requirement for obtaining cocaine was increased quasi-exponentially within-session, from FR1 to FR81 (see Methods). No response-contingent cue light was present in this experiment to avoid its behavioral confounding effects as shown above. Typically, with increasing FR values, responding for cocaine first increases, reaches a peak and then collapses to 0. The FR value corresponding to the peak of responding or maximum output (*O*max) represents the maximum price (or *P*max) that a rat is willing to pay to maintain cocaine self-administration (Fig. 6a). Both behavioral parameters are generally inter-correlated and also correlated with other measures of motivation, such as, for instance, the breakpoint in the classic progressive-ratio procedure. We measured *P*max and *O*max in the same rats before and after extinction of cocaine priming (F7,49 = 4.12, p < 0.01) (Fig. 6b). On the first post-extinction session of cocaine self-administration, rats were initially less motivated for the drug, as evidenced by a decrease in both *P*max (F5,35 = 3.29, p < 0.02) and *O*max (F5,35 = 2.73, p < 0.05) below the pre-extinction level (Fig. 6c,d). However, this decrease in drug motivation was short-lived. It recovered as early as the second post-extinction session onward. Similarly and not surprisingly, the number of cocaine injections per session - which directly depends on work output in the increasing FR procedure - also slightly decreased on the first post-extinction session but returned to normal afterward (F5,35 = 3.19, p < 0.02) (Fig. 6e) (see Fig. S5 for pre versus post-extinction patterns of cocaine self-administration).

**Figure 6.**
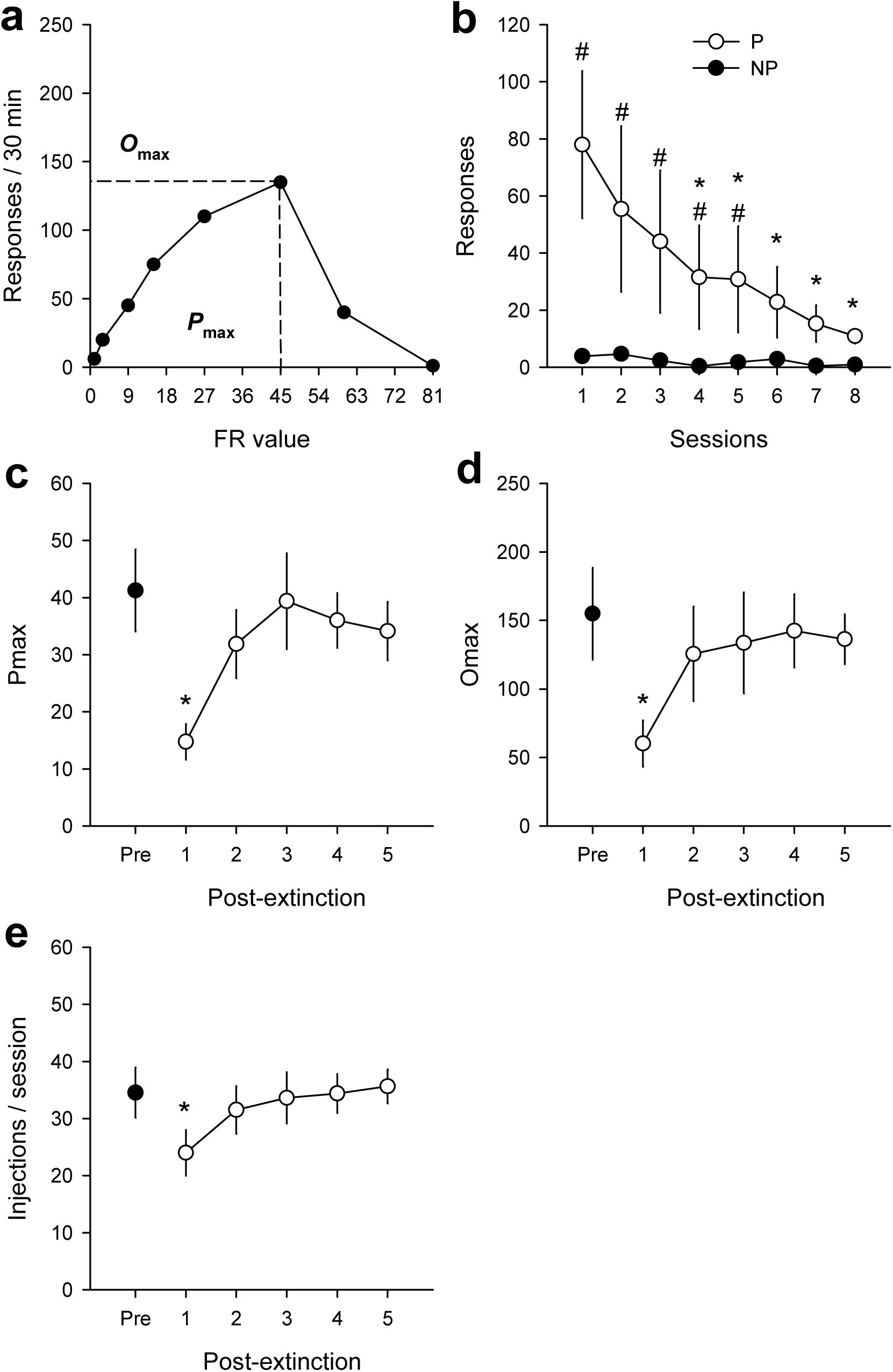
Post-extinction relapse to cocaine self-administration under a demanding FR procedure. (**a**) Graphic estimation of Pmax and Omax from a representative rat. In this case, the value of Omax was 135 responses/30 min and the value of Pmax was 45 responses/unit dose. (**b**) Extinction of the average effect of cocaine priming across sessions (mean ± s.e.m.). #, different from NP; *, different from session 1 (p < 0.01). (**c**) Pre-versus post-extinction Pmax values (mean ± s.e.m.). *, different from pre-extinction (p < 0.01). (**d**) Pre-versus post-extinction Omax values (mean ± s.e.m.). *, different from pre-extinction (p < 0.01). (**e**) Pre-versus post-extinction number of cocaine injections per session (mean ± s.e.m.). *, different from the last 2 post-extinction sessions (p < 0.01) but not from pre-extinction.

### Experiment 10: Neuronal correlates of extinction of cocaine priming

Since there was no evidence for a major impact of extinction of cocaine priming on relapse, we controlled for possible evidence for extinction-resistant neurobiological responses to cocaine in brain regions causally involved in cocaine-induced reinstatement. These regions include the anterior cingulate cortex, the prelimbic subdivision of the medial prefrontal cortex, and the core subdivision of the nucleus accumbens ^23, 24, 25^ (for exact anatomical coordinates, see ^51^). This experiment involved a total of 4 different subgroups of rats with different histories of extinction and different treatments during final testing (see Methods): subgroup V-V was repeatedly primed with vehicle during extinction and tested with vehicle; subgroup V-C was repeatedly primed with vehicle during extinction and tested with cocaine; subgroup C-V was repeatedly primed with cocaine during extinction and tested with vehicle; and finally, subgroup C-C was repeatedly primed with cocaine during extinction and tested with cocaine. As expected, during final testing, cocaine priming induced a large reinstatement effect in rats V-C, but not in rats C-C, compared to control rats that were tested with vehicle (i.e., rats V-V and C-V) (Group x Subgroup x Period interaction: F1,15 = 15.72, p < 0.01) (Fig. 7a). At the neuronal level, cocaine priming induced a large increase in Fos expressing-cells in all brain regions of interest in rats V-C compared to control rats (Group x Subgroup interaction: F1,15 values > 6.20, p values < 0.05) (Fig. 7b-d). These neuronal responses to cocaine priming were lost in rats C-C as they were not significantly different from those measured in control rats receiving vehicle (Figs. 7b-d). These results show that extinction of cocaine priming is associated with a shutdown of neuronal responses in brain regions critically involved in cocaine-induced reinstatement.

**Figure 7.**
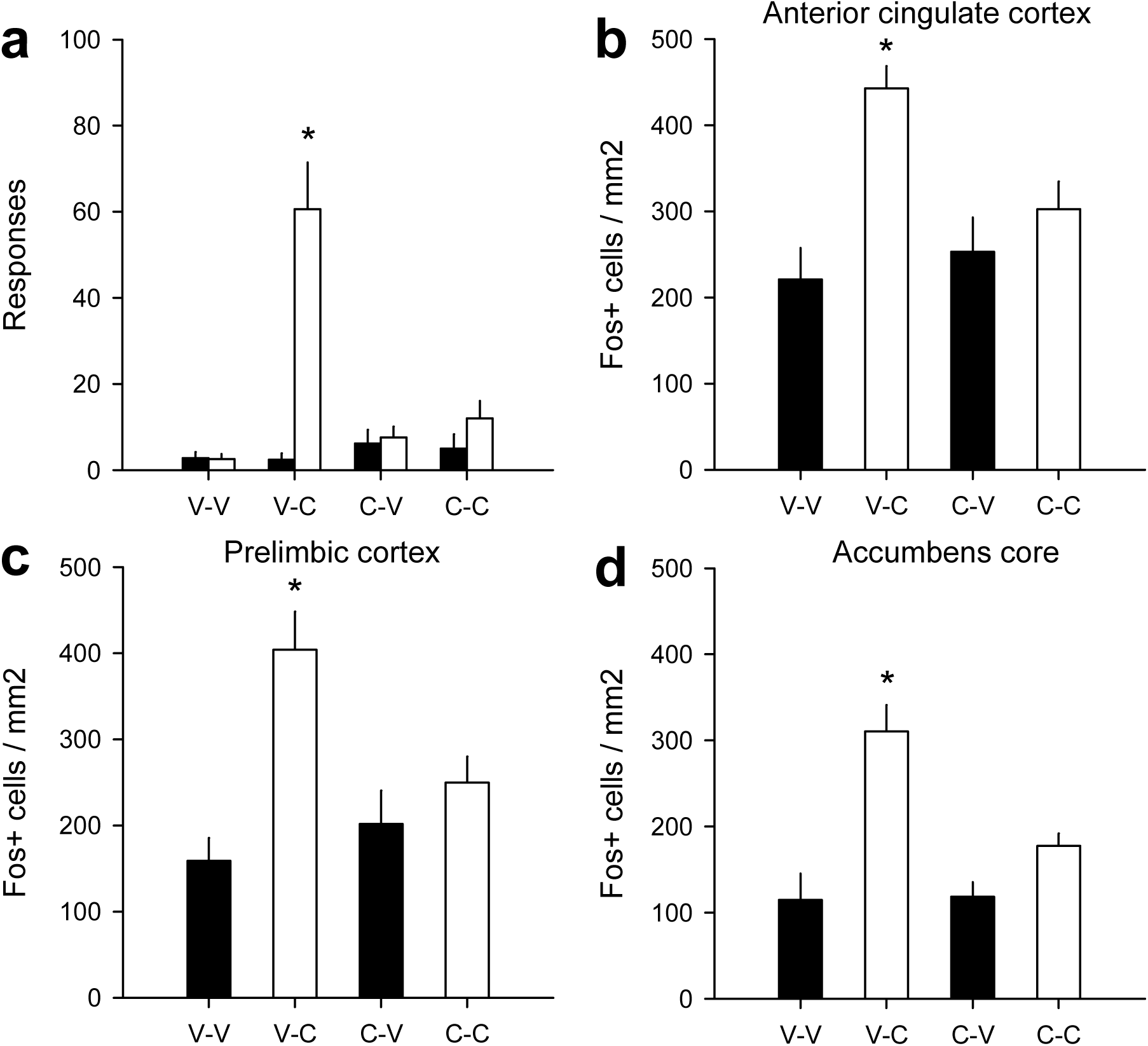
Neuronal correlates of extinction of cocaine priming. (**a**) Between-subject evidence for extinction of cocaine priming. Four subgroups of rats (n = 4-5) with different histories of extinction received either vehicle or cocaine during final testing. Subgroup V-V was repeatedly primed with vehicle during extinction and tested with vehicle; subgroup V-C was repeatedly primed with vehicle during extinction and tested with cocaine; subgroup C-V was repeatedly primed with cocaine during extinction and tested with vehicle; and finally, subgroup C-C was repeatedly primed with cocaine during extinction and tested with cocaine. The effect of each priming dose of vehicle or cocaine was compared to non-primed responding (NP) measured during the 45-min period preceding priming. *, different from all other conditions (p <0.01). Fos expression in the anterior cingulate cortex (**b**), the prelimbic cortex (**c**) and the nucleus accumbens core (**d**). Data are the mean (± s.e.m.) density of Fos-positive (Fos+) cells. *, different from control subgroups (p < 0.05).

Finally, *in vivo* electrophysiology revealed that extinction of the behavioral effects of cocaine on reinstatement of cocaine seeking (Fig. 8a) was paralleled in time by a dramatic extinction of its effects on neuronal activity in the prelimbic cortex (Fig. 8b). During the first extinction session, cocaine priming induced a robust reinstatement effect which was preceded by a rapid and prolonged increase in neuronal activity in the prelimbic cortex (Fig. 8b; see also, Fig. S6) that concerned most recorded neurons (i.e., 64 %). This finding provides direct *in vivo* evidence for the hypothesis that cocaine priming induced reinstatement of cocaine seeking via increased neuronal activity in the prelimbic cortex. It also validates, if needed, the interpretation of increased Fos expression in this brain region in terms of increased neuronal activity. With repeated priming, however, cocaine-induced increased neuronal activity in the prelimbic cortex progressively decreased to the NP level (Priming x Session: F4,116 = 8.62, p < 0.01) (Fig. 8b). This gradual neuronal adaptation paralleled in time the decrease in the reinstating effects of cocaine (Priming x Session: F4,8 = 11.04, p < 0.01) (Fig. 8a).

**Figure 8.**
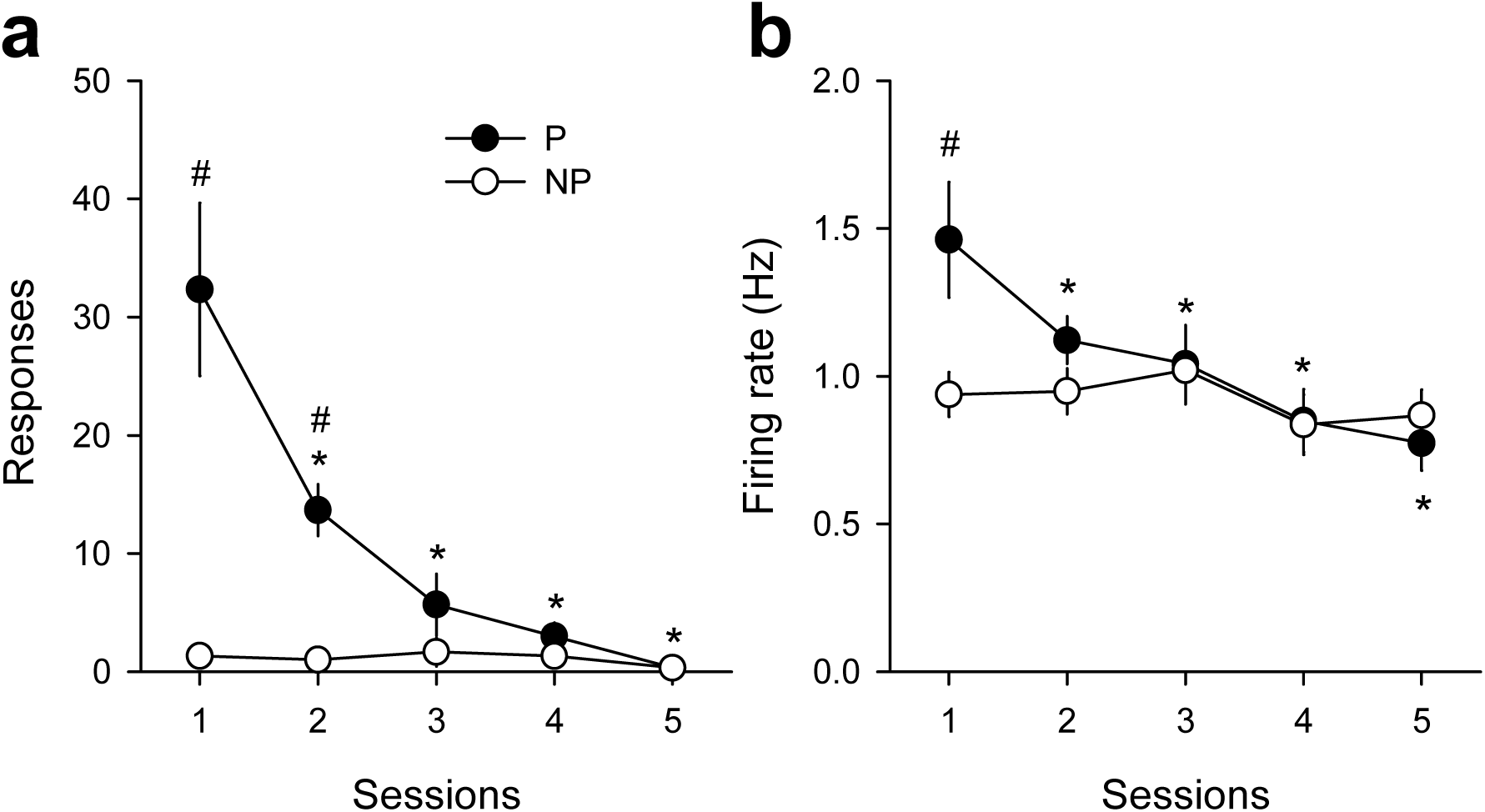
Parallel extinction of the behavioral and neuronal effects of cocaine priming. (**a**) Extinction of the effect of cocaine priming on non-reinforced responding across sessions (mean ± s.e.m.). The effect of cocaine priming (P) was compared to non-primed responding (NP) measured during the 45-min period preceding priming. (**b**) Mean firing rate (Hz) of neurons in the prelimbic cortex across repeated priming sessions (mean ± s.e.m.). The effect of cocaine priming (P) on prelimbic neuronal activity was compared to NP activity measured during the 45-min period preceding priming. #, different from NP; *, different from session 1 (p < 0.01).

Overall, even after extensive and complete extinction of cocaine priming, both at the behavioral and neural levels, there was no or only a modest and short-lived impact on relapse. As soon as rats found that cocaine was again available for self-administration during a session, they immediately returned (i.e., within one session) to their initial pre-extinction pattern of cocaine self-administration. Thus, the priming effect of cocaine seems to play no role in relapse vulnerability. These findings reveal, rather trivially, that the only way to prevent relapse would be to make rats stop responding entirely or at least sufficiently not to meet the operant response requirement for cocaine reinforcement. Clearly, this goal was not achieved in the present study, even during the first 90 min of non-primed responding. Despite evidence for complete extinction of cocaine priming, rats from experiments 3 and 5 that were tested for a total of 45 sessions of extinction continued nevertheless to exhibit a relatively high level of non-primed responding on the last 2 sessions of extinction (F17,289 = 13.30, p < 0.01) (Fig. 9a) and at a cumulative rate sufficient to meet rapidly several times the FR3 requirement for cocaine reinforcement (F17,289 = 11.21, p < 0.01; comparison with 3: all t17 values > 3.00, all p values < 0.01) (Fig. 9b). Clearly, this behavior insured that rats will rapidly detect cocaine when it was again available for self-administration after extinction of cocaine priming. Importantly, this behavior was not a side-effect of repeated cocaine priming because a similar behavior was also exhibited by control rats that were repeatedly primed with saline during extinction (Fig. S7: Responses: F17,170 = 12.27, p < 0.01; Cumulative responses: F17,170 = 15.10, p < 0.01; comparison with 3: all t10 values > 3.23, all p values < 0.01).

**Figure 9.**
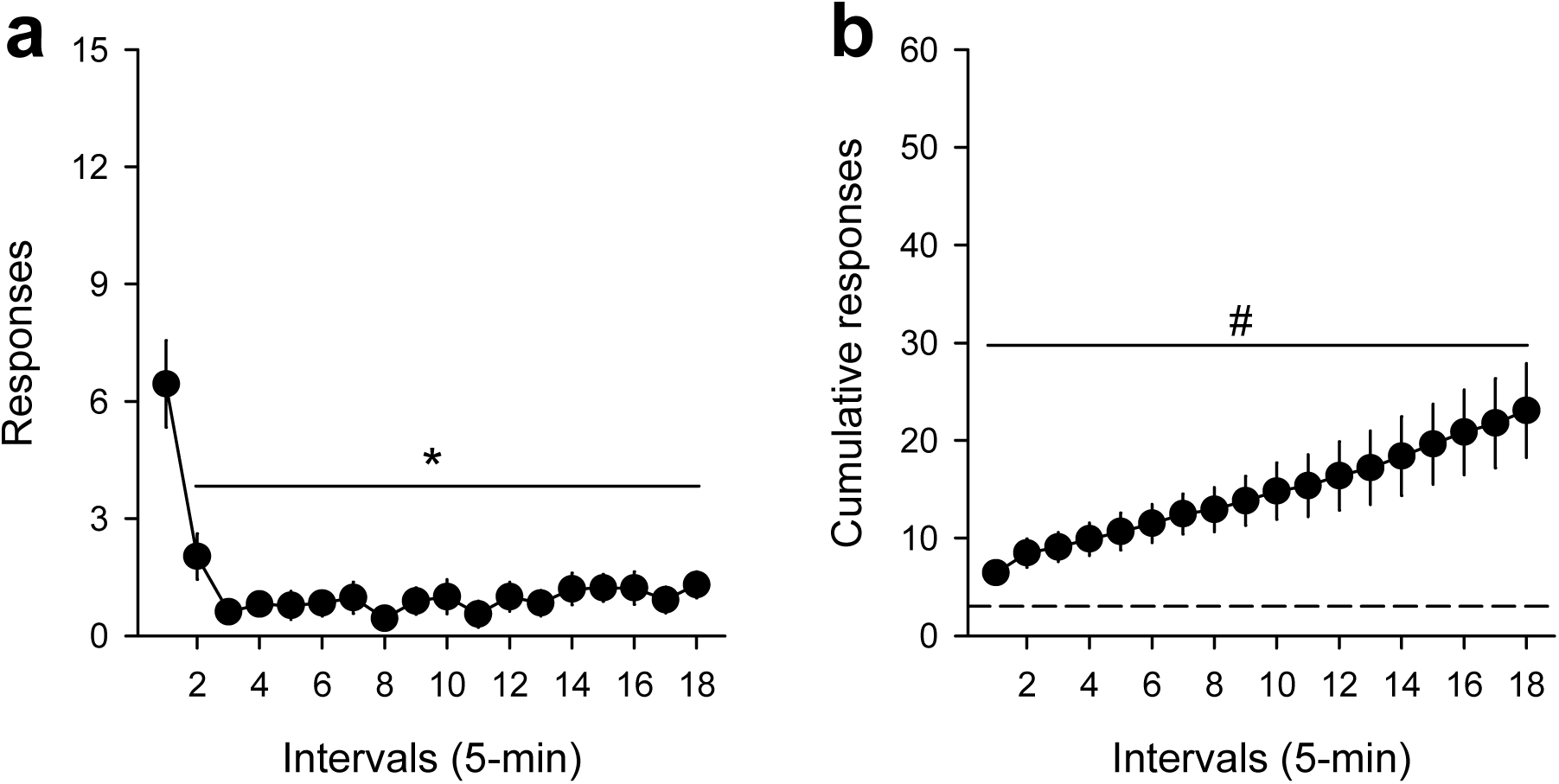
Continued non-primed responding after extinction of cocaine priming. (**a**) Within-session time course of non-primed responding after complete extinction of the priming effect of cocaine in rats that received a total of 45 sessions of extinction. *, different from the first 5 min (p < 0.01); (**b**) Cumulative non-primed responding in the same conditions as in (a). #, different from FR3 (p < 0.01).

## Discussion

The present study demonstrates that extinction of cocaine priming, whether administered contingently or non-contingently, has no major preventive effect on relapse. This lack of effectiveness could not be attributed to a residual effect of cocaine priming. When such residue was observed at the behavioral level, it was found to result from a cocaine potentiation of light seeking - a finding that confirms previous research with other psychostimulants ^19, 20, 21, 22^ - and not from cocaine seeking *per se*. In addition, at the neurobiological level, extinction of cocaine priming was associated with a loss of neuronal responses to cocaine in brain regions critically involved in cocaine-induced reinstatement, such as the prelimbic cortex and the nucleus accumbens core ^23, 24, 25^. Notably, in the prelimbic cortex, loss of neuronal responses to cocaine paralleled in time the extinction of its behavioral effects on reinstatement of cocaine seeking. In fact, even when extinction of cocaine priming was extensive and complete at the behavioral level, it had no major impact on subsequent relapse. An effect of extinction of cocaine priming was only observed when cocaine was available for self-administration under more demanding conditions. However, this effect was only modest and short-lived. In all relapse experiments conducted in the present study, once rats found that cocaine was again available for self-administration after extinction, they immediately recovered their initial pre-extinction self-administration behavior as if no extinction had occurred.

Overall, these findings demonstrate that the processes underlying cocaine-primed reinstatement do not play an essential role in vulnerability to relapse and that they are dissociable from the processes underlying the regulation of cocaine reward during self-administration. This behavioral dissociation was particularly evident in rats from experiment 3 that quickly relapsed after each block of extinction despite evidence for retention of extinction of cocaine priming across blocks. During the last blocks, the same individual rats no longer responded to the priming effects of cocaine but nevertheless continued to self-administer cocaine at the same rate and according to the same pattern as before extinction. Lack of responses to the priming effects of cocaine co-existed, within the same individuals, with an unchanged cocaine self-administration behavior. This within-subject behavioral dissociation confirms several between-subject neurobiological dissociation studies showing that neural interventions that reduce or block cocaine-primed reinstatement have no effect on cocaine self-administration ^26, 27, 28, 29, 30, 31, 32, 33, 34^. This dissociation may not be entirely surprising, however. It probably indicates that the motivational processes underlying the regulation of cocaine reward during self-administration is largely independent from the Pavlovian associative processes hypothesized to be involved in the priming effects of cocaine (see Introduction). To make a simple analogy: as extinction of food priming in hungry rats would not be expected to eliminate hunger, extinction of cocaine priming should perhaps also not be expected to eliminate the motivation to use cocaine ^35, 36^. For instance, it is likely that the hedonic set point mechanism that normally underlies and motivates regulated cocaine self-administration will largely remain intact after extinction of the conditioned interoceptive cues of cocaine, or any other conditioned cues for that matter ^37, 38^. Regardless of the significance of this dissociation, the present failure to block relapse after complete extinction of cocaine priming indicates that contrary to our initial hope, this extinction strategy may not represent a promising strategy to prevent relapse in people with cocaine addiction ^11^. Whether this conclusion can also be extrapolated to other strategies that also target the priming effects of cocaine or related cues is currently unknown, mainly because, apart from rare exceptions ^39^, the effects of these strategies on relapse prevention have not been tested yet as directly as here.

We were able to trace the origin of our failure to prevent relapse to an extinction-resistant form of operant responding that systematically occurred at the beginning of each session and that, though relatively weak, was nevertheless sufficiently elevated to meet the previously learned response requirement for cocaine reinforcement (i.e., FR3 in the present study) (see Fig. 9). Obviously, this persistent behavior was a precondition for rapid relapse because it allowed rats to rapidly and systematically detect the post-extinction return of cocaine reinforcement in the present study. Though the mechanisms underlying this extinction-resistant behavior remain to be fully elucidated, it may merely reflect rats’ attempt - in the absence of other predictive information - to check whether the availability of cocaine reinforcement in a given session has changed or not from the previous session. In theory, though lack of cocaine reinforcement over repeated extinction sessions should lead rats to expect lack of reinforcement in a current session, there is nevertheless always a possibility of change in drug availability that would be missed if rats stopped totally to respond. In wild habitats, when a particular source of reinforcement is depleted, it is generally not permanently. It may eventually be replenished (e.g., a patch with earth worms may be replenished by the arrival of new worms). In other words, things may and do change ^40, 41^. Thus, for rats that are still motivated to take cocaine (and that have no access to other competing pursuits), a cost-effective, adaptive strategy to avoid missing an eventual post-extinction change in cocaine availability is merely to continue to respond at least at the beginning of each session and at a sufficiently high level. This strategy is cost-effective in the sense that it is relatively effortless as it only requires rats to emit few initial responses to obtain the relevant information. Indeed, if responding is not reinforced at the beginning of the session, then this is confirmation that, as expected from previous extinction sessions, cocaine will not be available, thereby prompting rats to stop responding for the remaining of the session. Though the extinction-resistant behavior reported here is entirely consistent with the above interpretation in terms of information seeking in cocaine-motivated rats, further research will nevertheless be needed to fully demonstrate its validity.

Regardless of the underlying psychological mechanisms, however, it is clear that this persistent, extinction-resistant behavior is responsible for our systematic failure to prevent relapse in the present study. Thus, eradicating this behavior seems to be a pre-requisite for relapse prevention in animal models of cocaine addiction. To our knowledge, this outcome has so far never been achieved, including in recent promising studies that also succeeded to reduce the priming effects of cocaine ^39^. In fact, a quick perusal of the relevant literature reveals that rats never stop entirely to respond, even after prolonged extinction (e.g., all articles on cocaine-induced reinstatement cited in the References). As reported here, they generally continue to respond at a weak, albeit largely sufficient level (i.e., between 10 and 20 responses) to detect cocaine reinforcement if it happened to be again available after extinction. Extinction-resistant responding is thus one of the most reproducible, albeit largely overlooked, behavioral outcomes of drug reinstatement and extinction studies. Of course, this persistent behavior may not be entirely surprising if one considers that it is relatively effortless compared to its potentially large payoff (i.e., discovery of return of cocaine availability) and that it is not in competition with other rewarding pursuits ^42^. Clearly, future research on relapse prevention should focus on the different factors of such persistent, extinction-resistant behavior.

Finally, the present study also provides some new insights about the neural mechanisms of cocaine-primed reinstatement of drug seeking. Previous research has amply demonstrated that the dorsomedial prefrontal cortex, including its anterior cingulate and prelimbic subdivisions, plays a causal role in cocaine-primed reinstatement of cocaine seeking, notably through its top-down projections to the core of the nucleus accumbens ^23, 24, 25, 43^. The present study is generally consistent with this conclusion. In addition, the fact that cocaine no longer activated this top-down prefrontal-accumbens pathway after repeated priming suggests that its recruitment is probably a conditioned response to the interoceptive stimuli of cocaine priming, as hypothesized here. One important virtue of this interpretation is that it explains cocaine-primed reinstatement as an instance of a more general mechanism of conditioned cue-or context-induced cocaine seeking ^7, 11, 44^. In all cases, the starting point is the perception of a drug-predictive cue or context that leads rats to falsely expect, at least initially, the return of cocaine reinforcement after operant extinction. The only difference would be the interoceptive *versus* exteroceptive sensory modality of the drug-predictive stimuli and, perhaps, the different subcomponents of the prefrontal-accumbens pathway that they probably recruit. What the conditioned interoceptive stimuli of cocaine priming are exactly, where they are processed in the brain, and how the output of this processing leads to activation of the prefrontal-accumbens pathway that controls reinstatement of cocaine seeking remain important unanswered questions for future research ^13^.

In conclusion, the present study reveals that complete extinction of cocaine priming does not prevent relapse in rats. This failure to prevent relapse was mainly due to the existence of a persistent, extinction-resistant form of operant behavior that probably reflects a continued motivation to seek cocaine. Future research should be reoriented to focus on this specific behavior both as a potential marker for persistent vulnerability to relapse and as a potential target for the development of potential relapse prevention interventions. The failure to target and eradicate this behavior may explain, at least partly, why preclinical research has so far been unable to deliver clinically-effective relapse prevention interventions in cocaine addiction ^45, 46^.

## Methods

### Animals and Housing

A total of 171 adult male Wistar rats (225-250 g at the beginning of experiments, Charles River, Lyon, France) were used. Rats were housed in groups of 2 and were maintained in a light-(reverse light-dark cycle), humidity-(60 ± 20%) and temperature-controlled vivarium (21 ± 2°C). All behavioral testing occurred during the dark phase of the light-dark cycle. Food and water were freely available in the home cages throughout the duration of the experiment. Home cages were enriched with a nylon gnawing bone and a cardboard tunnel (Plexx BV, The Netherlands). 49 rats did not complete the experiment because of catheter failure, infection or a failure to meet the criteria for acquisition of cocaine self-administration (> 12 injections per 3-h session), thereby leaving 122 rats for final analysis.

All experiments were carried out in accordance with institutional and international standards of care and use of laboratory animals [UK Animals (Scientific Procedures) Act, 1986; and associated guidelines; the European Communities Council Directive (2010/63/UE, 22 September 2010) and the French Directives concerning the use of laboratory animals (décret 2013-118, 1 February 2013)]. The animal facility has been approved by the Committee of the Veterinary Services Gironde, agreement number A33-063-922.

### Surgery

Rats were surgically prepared with an indwelling silastic catheter in the right jugular vein under deep anesthesia. Behavioral testing commenced at least 7 days after surgery. Additional information about surgery and post-operative care can be found elsewhere ^47^.

### Apparatus

Fourteen identical operant chambers (30 x 40 x 36 cm) were used for all behavioral testing and training (Imétronic, Pessac, France). Each chamber was equipped with two retractable metal levers on opposite panels of the chamber, and a corresponding white cue light was positioned above each lever. There was also an array of 5 identical white cue lights on the roof of each chamber, 4 in the roof corners and one in the middle. Finally, each cage was also equipped with two transversal photocell beams for measuring cage cross overs as a proxy for horizontal locomotion. Additional information about the apparatus can be found elsewhere ^48^.

### Drugs

Cocaine hydrochloride (Coopération Pharmaceutique Française, Melun, France) was dissolved in 0.9% NaCl, filtered through a syringe filter (0.22 μm) and stored at room temperature. Drug doses are expressed as the weight of the salt.

### Initial cocaine self-administration training

In all experiments, rats were first habituated during 2 3-h daily sessions to the experimental chambers. During habituation, no lever or light cue was presented, and rats were allowed to move freely and explore the cage. After habituation, rats were progressively trained to press a lever to self-administer cocaine intravenously (0.25 mg per injection) under a final fixed-ratio (FR) 3 schedule of reinforcement during 3-h daily sessions. The number of self-administration sessions before extinction of cocaine priming ranged from 24 to 64 depending on the experiment (see below). All self-administration sessions began with extension of the operant lever and ended with its retraction after 3 h. No inactive lever was used in the present study. Intravenous delivery of cocaine began immediately after completion of the operant response requirement and lasted about 4 s. It was generally accompanied by illumination of the light cue above the lever for 20 s, except when indicated (see below). Responses during the light cue were recorded but had no programmed consequence. All self-administration sessions were run 6-7 days a week.

### Within-session multiple priming procedure for extinction of cocaine priming

Extinction of cocaine priming began 1 day after the last session of cocaine self-administration and was induced using a multiple drug priming variant of a within-session extinction-reinstatement procedure described previously ^11, 49, 50^. Briefly, each session lasted 225 min and consisted of an initial 90-min period of extinction of operant responding during which completion of the FR3 continued to turn on the 20-s light cue above the operant lever (except when otherwise indicated) but was no longer reinforced by cocaine. After this initial period of 90 min, rats continued to be tested under operant extinction but received 3 1-mg priming doses of cocaine (P1 to P3) spaced 45 min apart. Each priming dose was generally administered non-contingently, except when otherwise mentioned (see experiment 2 below). The dose of 1 mg was previously shown to induce a maximal priming effect on reinstatement of cocaine seeking in a previous study ^11, 49^. Three priming doses of cocaine per session were used in an attempt to promote a rapid and complete extinction of cocaine priming. Cocaine-induced reinstatement of operant responding was measured during the 45-min periods following P1, P2 and P3, and compared to non-primed responding (called NP) measured during the 45-min period preceding P1. Depending on the experiment, the number of extinction sessions before relapse testing ranged from 5 to 45 (see below).

### Post-extinction relapse to cocaine self-administration

One day after the last session of extinction of cocaine priming, cocaine was made again available for self-administration during 5-6 consecutive sessions under the same operant conditions as before extinction. No explicit cue announced post-extinction cocaine availability, except for the availability of cocaine reinforcement itself upon operant responding. In all experiments, the impact of extinction of cocaine priming on relapse was assessed using a within-subject comparison of the pre-versus post-extinction patterns and rates of cocaine self-administration.

### Specific experiments

Several separate experiments were conducted to measure the impact of extinction of cocaine priming on relapse to cocaine self-administration.

#### Experiment 1: Effects of extinction of non-contingent priming cocaine on relapse

This initial experiment was conducted as a pilot study in a group of rats (n = 20) from a previous, unrelated experiment. Before being exposed to repeated sessions of extinction of cocaine priming, these rats had been extensively tested for cocaine self-administration under different conditions. In total, they self-administered cocaine during 64 daily sessions of at least 3h each whose 37 3-h sessions under a FR3 schedule of reinforcement. Extinction of cocaine priming was induced during 5 daily sessions using the triple priming procedure described above. Then one day after the last session of extinction, rats were tested for relapse to cocaine self-administration during 5 days as described above.

#### Experiment 2: Effects of extinction of response-contingent cocaine priming on relapse

Despite evidence for a large extinction of cocaine priming in experiment 1, there was apparently no significant impact on subsequent relapse to cocaine self-administration. Experiment 2 tested whether this lack of effect is attributable to the non-contingent nature of cocaine priming. To test this, an independent group of rats (n = 28) previously trained to self-administer cocaine during 24 sessions, as described above, were tested under a modified multiple priming procedure where all 3 priming doses of cocaine were under rats’ control. Specifically, they were delivered only if rats pressed one time on the lever after onset of the relevant period. Importantly, however, to avoid excessive session duration in case of delayed responding or omission, a maximum latency of 15 min was allowed for cocaine self-priming. If self-priming was initiated within this delay, this triggered a 45-min period during which no other priming was delivered and operant responding was non-reinforced. If rats failed to respond within the maximum delay, no priming was administered and this began the next period of opportunity for self-priming. There was a total of 3 opportunity periods per session. Finally, to equate conditions, rats were also required to self-initiate the 45-min NP period that precedes the first self-priming. Note that under this self-priming procedure, it was possible for a rat to fail to self-administer all 3 primings (see Results). Rats were tested under this procedure during 6 sessions before being tested for relapse.

#### Experiment 3: Effects of extensive extinction of cocaine priming on relapse

This experiment sought to measure the effects of more extensive extinction of cocaine priming on relapse to cocaine self-administration. An independent group of rats (n = 18), previously trained to self-administer cocaine during 24 sessions, were tested during 24 sessions of extinction of cocaine priming. These sessions of extinction were organized into 4 blocks of 6 sessions that alternated with 4 blocks of 6 sessions of relapse to cocaine self-administration. This design allowed us to follow any change in vulnerability to relapse with increased sessions of extinction.

#### Experiment 4: Effects of saline substitution on residual cocaine priming after extinction

In all experiments described above, there remained a small, residual priming effect of cocaine, even after extensive extinction testing. This experiment sought to measure the contribution of the intravenous injection procedure itself to this residual effect. This was particularly warranted in the present study because no intravenous injection was administered during the control NP period. To this end, a separate group of rats (n = 13), trained to self-administer cocaine during 35 sessions, were tested during 5 sessions of extinction of cocaine priming until evidence for a residual effect of cocaine emerged. Then, all else being equal, cocaine was replaced with 0.9% saline - its vehicle. This saline substitution session was replicated 3 times between regular sessions of cocaine reinstatement.

#### Experiment 5: Effects of light cue conditions on residual cocaine priming after extinction

As explained above, completion of the FR3 continued to be signaled by a 20-s light cue above the operant lever during extinction sessions. Though this cue should have lost most of its conditioned effects after extensive extinction of both operant responding and cocaine priming, its continuous response-contingent presentation may nevertheless contribute to the residual effect of cocaine priming through either some extinction-resistant conditioned reinforcing effects or other unconditioned reinforcing mechanisms. To test this, rats from experiment 3 (n = 18) received additional sessions of extinction under the following 3 light cue conditions: 1) light cue available (LC, standard extinction condition): the 20-s light cue previously paired with cocaine is presented above the operant lever upon completion of the FR3; 2) no light cue is available (NLC): completion of the FR3 has no programmed consequence throughout the session; and, finally, 3) house light cue available (HLC): a novel light cue - an array of 5 house light cues from the cage roof - never paired with cocaine is presented upon completion of the FR3. Each cue condition was tested during at least 4 consecutive sessions in the following, partially mixed, order: LC condition (9 sessions), NLC condition (4 sessions), HLC condition (4 sessions), NLC condition (2 sessions) and, finally, LC condition (2 sessions).

#### Experiment 6: Effects of cocaine on operant responding for a neutral light cue after extinction of cocaine priming

A separate group of rats (n = 13) was first trained to self-administer cocaine with the response-contingent light cue during 17 sessions, as described above. Then the response-contingent light cue was suspended for an additional 13 sessions of self-administration and throughout extinction testing (NLC condition). After 8 sessions of extinction under the NLC condition, the house light cue (HLC) described in experiment 5 was introduced and made contingent upon completion of the FR3 during a final session of extinction. The priming effect of cocaine under the HLC condition was compared to that measured during the last 3 sessions of extinction under the NLC condition.

#### Experiment 7: Effects of cocaine on operant responding for a neutral light cue in naïve rats

Experiment 7 sought to measure the direct effects of cocaine on operant responding for a 20-s light cue in initially operant-and drug-naïve rats. After habituation to the experimental chambers, as described above, rats (n = 28) were given access to one lever during 2 sessions of 225 min under two different light cue conditions. In one session, FR1 responding on the lever had no programmed consequence (NLC session); in the other session, FR1 responding turned on a 20-s light cue above the operant lever (LC session). Each session began with presentation of the lever and ended with its retraction. During both sessions, after an initial 45 min with no drug injection, rats received 4 increasing doses of intravenous cocaine (0, 0.25, 0.5 and 1 mg per injection) each spaced 45 min apart. This procedure was designed to be as close as possible to the standard triple priming procedure used in the previous experiments. After this short experiment, the same rats were then used as subjects in experiment 2 described above.

#### Experiment 8: Effects of extensive and complete extinction of cocaine priming on relapse

This experiment was a follow-up of experiments 3 and 5. After a total of 45 sessions of extinction and evidence of complete extinction of cocaine priming, rats (n = 18) were tested for relapse to cocaine self-administration during 2 consecutive sessions as described above.

#### Experiment 9: Effects of extinction of cocaine priming on relapse under a more demanding procedure of cocaine self-administration

In all previous experiments, a relatively moderate level of effort (i.e., FR3) was required to obtain cocaine during both maintenance and relapse to cocaine self-administration. This experiment tested rats under a more behaviorally demanding procedure of cocaine self-administration before and after extinction of cocaine priming. To this end, we used a group of rats (n = 8) from an unrelated experiment. In total, these rats self-administered cocaine during 52 sessions with no light cue, except for the first 16 sessions. No light cue was used to avoid the associated bias uncovered in the 3 preceding experiments. Then they were trained under a within-session increasing FR procedure ^16^. All other procedural details for cocaine self-administration were as described above. Briefly, after an initial 15-min period of cocaine loading under a FR1 schedule, the FR value was increased quasi-exponentially from 1 to 81 every 30 min (i.e., 1, 3, 9, 15, 27, 45 and 81). Rats were tested under this increasing FR procedure during 6 consecutive sessions before and 5 consecutive sessions after 8 sessions of extinction of cocaine priming.

#### Experiment 10: Neuronal correlates of extinction of cocaine priming

This experiment used c-Fos immunohistochemistry to check whether extinction of cocaine priming was associated with changes in neuronal responses to cocaine priming in brain regions critically involved in cocaine-induced reinstatement (see below). Rats (n = 19) were first trained to self-administer cocaine during 30 sessions, as described in the General Behavioral Procedures (i.e., TO cue present). Then they were distributed into 2 balanced groups (i.e., same levels of cocaine intake) for extinction testing. One control group was primed with vehicle (Group V) while the other group was primed with 1 mg of cocaine (Group C). Extinction testing was identical to previous experiments except that only one non-contingent priming was administered per session. This was done to minimize the difference in cocaine exposure between groups V and C before the final challenge session. After evidence for stable extinction of cocaine priming in group C, each group was further subdivided into two subgroups during a final testing session, one primed with vehicle, the other with 1 mg of cocaine. Thus, there was a total of 4 different subgroups (n = 4-5): subgroup V-V (repeatedly primed and challenged with vehicle), subgroup V-C (repeatedly primed with vehicle and challenged with cocaine), subgroup C-V (repeatedly primed with cocaine and challenged with vehicle), and subgroup C-C (repeatedly primed and challenged with cocaine). Subgroup V-V controlled for the effects of cocaine priming in subgroups V-C and C-C and subgroup C-V controlled for any eventual carry-over effects due to previous repeated cocaine priming. At the end of the challenge session (i.e., 45 min after priming), all rats were anesthetized and perfused transcardially with 4% formaldehyde. Then their brains were extracted, post-fixed, sliced, and processed for Fos immunohistochemistry analysis as described in details elsewhere ^50, 51^. Density of Fos-expressing cells (cells / mm2) was measured by a blind observer in both hemispheres of each rat and averaged from 3 sections covering the anterior-posterior (AP) extent of each region of interest. Brain regions of interest were defined here as regions previously reported to be consistently involved in cocaine-induced reinstatement. These regions include the anterior cingulate cortex, the prelimbic subdivision of the medial prefrontal cortex, and the core subdivision of the nucleus accumbens ^23, 24, 25^ (for exact anatomical coordinates, see ^51^). We plan to eventually report a more comprehensive and fine-grained Fos mapping of the neuronal correlates of extinction of cocaine priming in a separate article. Here our goal was limited to confirming that extinction of cocaine priming was also neurobiologically effective.

Finally, to assess whether there was a temporal relationship between behavioral and neuronal changes during extinction of cocaine priming, we also used *in vivo* electrophysiology to record single-unit activity in the prelimbic subdivision of the medial prefrontal cortex in an independent subset of rats (n = 3). Cocaine is thought to reinstate cocaine seeking via activation of prelimbic cortex neurons that project to the core of the nucleus accumbens ^52^. Here we sought to directly demonstrate this activation *in vivo* and follow its evolution during repeated cocaine priming. Rats were first trained to self-administer cocaine during 44 sessions under a FR1 schedule of reinforcement, as described above. Then they were implanted unilaterally under isoflurane anesthesia with arrays of 16 teflon-coated stainless steel microwires (MicroProbes Inc, Gaithersburg, MD) in the prelimbic cortex [AP: + 2.5 to + 4.2 mm, ML: 0.5 to 1.5 mm, and DV: - 4.0 mm relative to skull level]. Ten days after surgery, rats were retrained to self-administer cocaine for 9 additional FR1 sessions before being tested for extinction of cocaine priming. Extinction testing was identical to previous experiments except that only one non-contingent drug priming (1 mg, i.v.) was administered per session. In addition, no response-contingent light cue was present throughout extinction testing to rule out its possible influence on recorded neuronal activity. Electrophysiological recordings were conducted during each daily extinction session. Within each session, prelimbic neuronal firing activity (in Hz) measured 45 min after cocaine priming (P) was compared to that measured during the preceding non-primed period of 45 min (NP). Neurons were recorded using commercial hardware and software (OmniPlex, Plexon, Inc, Dallas, TX) and classified into putative interneurons (n = 30) and pyramidal neurons (n = 160) according to the waveform spike width and average firing rate, as previously described ^53^. Due to the small number of interneurons, data analysis was exclusively focused on putative pyramidal neurons. To count the number of individual neurons that changed their firing rate in response to cocaine priming during each session, we compared firing rate measured 25 min before and 25 min after cocaine priming using the Wilcoxon test.

### Data Analysis

All data were subjected to relevant repeated measures ANOVAs, followed by Tukey post hoc tests where relevant. Comparisons with a fixed theoretical level (e.g., 3 responses) were conducted using one sample t-tests. Statistical analyses were run using Statistica, version 7.1 (Statsoft Inc., Maisons-Alfort, France).

## Acknowledgements

This research was supported by funding from the Centre National de la Recherche Scientifique (CNRS), the Agence Nationale de la Recherche (ANR-2010-BLAN-1404-01), the Université de Bordeaux, the Conseil Regional d’Aquitaine (CRA11004375/11004699), and the Labex BRAIN.

## Author contributions

S.H.A. conceived the project. P.G., K.G., S.N., A.D. and C.V-M. performed the experiments. P.G., K.G., S.N. and S.H.A. designed the experiments and analyzed the data. S.H.A. wrote the paper with the help of all of the other authors. All authors critically reviewed content and approved final version for publication.

## Additional information

**Supplementary information** accompanies this paper

**Competing financial interests:** the authors declare no competing financial interests.

